# Protein-Ligand Binding Site Prediction and *De Novo* Ligand Generation from Cryo-EM Maps

**DOI:** 10.1101/2023.11.16.567458

**Authors:** Chunyang Lu, Kaustav Mitra, Kiran Mitra, Hanze Meng, Shane Thomas Rich-New, Fengbin Wang, Dong Si

**Affiliations:** Division of Computing and Software Systems, University of Washington, 18115 Campus Way NE, Bothell, 98011, WA, United States; Department of Computer Science, Duke University, 308 Research Drive, Durham, 27708, NC, United States; Department of Biochemistry and Molecular Genetics, University of Alabama at Birmingham, 1720 2nd Ave S, Birmingham, 35233, AL, United States

**Keywords:** Protein-Ligand Binding Site Prediction, Machine Learning, Bioinformatics, Drug-Like Molecule Generation, Cryo-EM, Drug Design

## Abstract

Identification of protein-ligand binding sites is one of the most challenging tasks in drug discovery and design. Recent advances in machine learning community, particularly deep learning, inspired considerable research into deep learning-based methods for protein ligand binding site prediction (PLBP) and have achieved promising results. However one limitation of these methods and models is that they are developed and evaluated using conventional databases consisting of relatively small protein structures determined from X-ray crystallography and NMR with high resolution. When current PLBP methods are directly applied to large protein complexes, they either fail or produce very poor results. Given the increasing popularity of protein structures determined from Cryo-EM maps, which usually capture large protein complexes at relatively low resolution, there is a strong need to apply PLBP methods to Cryo-EM maps. For this purpose, we created a novel database (EMD-Ligand Dataset) consisting of Cryo-EM maps and information about their ligand-binding partners. Our study of several state-of-the-art PLBP methods show that even though they perform reasonably well on conventional databases, they do poorly on the novel EMD-Ligand Dataset, which calls for more research and work to be done by the community. As current methods for determining protein structures experimentally from Cryo-EM maps are time-consuming and expensive, we integrated PLBP methods with DeepTracer, an automatic and fast, de novo Cryo-EM protein structure modeling method, and established an end-to-end system capable of predicting ligand binding sites directly from Cryo-EM maps. Additionally, we leveraged an existing ligand generator to generate drug-like ligands from our predicted ligand-binding pockets and demonstrated its effectiveness.

## 1 Background

### 1.1 Motivation

Proteins perform crucial biological functions through their interactions with binding partners (ligands). Interactions that are critical for various pathologies have frequently been the subject of study in drug design, however, designing drugs that bind specifically to a protein’s unique three-dimensional structure is often challenging and can be costly on several fronts including time, chemical resources, and financially. To mitigate these factors, computational prediction of protein-ligand interactions has recently been integrated into the modern drug discovery field. Most recently, due to an abundance of experimental structural data, machine learning and artificial intelligence have been applied to investigate protein-ligand interactions. Prediction of proteinligand binding sites is essential for structure-based drug design, which may assist in predicting potential therapeutic side effects. Protein-ligand bindings are typically highly specific intermolecular interactions that occur at binding pockets on proteins.

There is a limitation in current popular deep-learning based protein-ligand binding site prediction (PLBP) methods such as Kalasanty [1], PUResNet [2] and DeepSurf [3]. These methods are developed and evaluated on public datasets like ScPDB [4] and PDBbind [5], which are mainly comprised of small protein complexes determined by X-ray crystallography and Nuclear Magnetic Resonance (NMR) with high resolution, typically *<*3Å. When directly applied to large protein complexes, these PLBP methods either fail or produce very poor results due to their limited experience with larger protein sizes and greater numbers of atoms. Given the increasing popularity of protein structures determined by Cryo-EM maps, which are usually images of large protein complexes at relatively low resolution, there is a strong need to study PLBP methods on proteins from Cryo-EM maps. To our knowledge, there is no published database explicitly designed for predicting ligand binding sites on proteins with Cryo-EM maps, which motivated us to create a novel database for this research: the EMD-Ligand Dataset. The Cryo-EM maps used in this database were retrieved from the Protein Data Bank (PDB) linked to EMDB, which is the database that has the Cryo-EM maps. This database enabled us to test the possibility of extending the functionality of DeepTracer [6], allowing for the prediction of protein-ligand binding sites directly from Cryo-EM maps without experimentally-determined protein 3D structures, as well as ligand generation using protein structures determined by DeepTracer.

### 1.2 Previous Protein-Ligand Interaction Prediction Methods

There has been considerable research in applying deep learning-based methods to predict ligand binding sites. For example, the SURFNET [6] architecture successfully predicted the ligand-binding site as the largest pocket in over 80% of all tested cases. Additionally, other methods such as MaSIF [7], DeepCSeqSite [8], DELIA [9], Kalasanty [1], DeepSurf [3], and PUResNet [2] have been developed to predict binding sites. MaSIF exploits geometric deep learning to learn interaction fingerprints in protein molecular surfaces. DeepCSeqSite is a sequence-based method using a deep convolutional neural network (dCNN) to predict protein-ligand binding residues. DELIA is a hybrid deep neural network integrating a CNN with a bidirectional long short-term memory network (BiLSTM) to mobilize 1D sequence feature vectors and 2D distance metrics. It improves the data imbalance between binding and nonbinding residues using the over-sampling in minibatch, random under-sampling and stacking ensemble strategies. Kalasanty is a 3D convolutional neural network based on U-Net architecture [10]. DeepSurf is based on deep learning architecture which mobilizes surface-based representation (implementation of 3D voxelized grids) along with state-of-the-art deep learning architectures to predict potential druggable sites on proteins. PUResNet implemented a deep ResNet as the backbone of the network, which is given as input a 3D protein structure and outputs the ligandability of each voxel. It also used a unique training data cleaning process.

Current PLBP methods rely on publicly available datasets like ScPDB [4] and PDBbind [5] to train and evaluate their models. However, these datasets mainly comprise smaller protein complexes determined by X-ray crystallography and NMR with high resolution. The “resolution revolution” of Cryo-EM that began in 2014 has enabled the imaging of an increasing number of structures using the technique, resulting in an increasing proportion of protein structures in the PDB Database[11] derived from Cryo-EM maps. This increase and the lack of PLBP methods utilizing Cryo-EM maps represents an unmet need in the study of protein-ligand interactions, drug discovery, and drug design. Further, Cryo-EM is an indispensable tool for studying the structures and functions of macromolecular assemblies and provides another basis for the design of novel therapeutic drugs.

In addition to the protein and ligand binding site prediction methods discussed earlier, there are several other related methods in the field. For instance, SSnet utilizes a 1D representation based on the curvature and torsion of the protein backbone. This method represents the protein using the Cα carbons of the backbone, which clearly defines a recognizable structure for peptides and proteins alike and is very useful for categorization [12]. Ligands are represented using SMILES strings, which represent various chemical bonds and orientations present in the molecule. Another approach, utilizing in silico chemogenomic informatics (which is based on mining the chemical space of the molecules in question) predicts the interactions between small molecules and proteins, which is crucial for understanding many biological processes and plays a critical role in drug discovery since this approach can identify and create a selectivity profile of similar proteins (homologues) in order to prevent side effects [13]. Docking approaches and quantitative prediction of binding affinity using regression methods in the joint space are also commonly used when the target 3D structure is available. SIGN [14] is a structure-aware interactive graph neural network that comprises polar-inspired graph attention layers (PGAL) and pairwise interactive pooling (PiPool) and is used for prediction of binding affinities as well as per-atom structural and interaction information. DeepAffinity [15] is a deep learning technique based on a recurrent neural network that represents a protein with amino acid sequences and a molecule with SMILES. This method embeds 3D structural information of a proteinligand complex in an adjacency matrix, devises a distance-aware attention algorithm to differentiate various types of intermolecular interactions, and introduces a variant of graph neural networks suitable for learning protein-ligand interactions. DeepDock [16] is a geometric deep learning-based method that extracts features from the input data to identify key ligand-target interactions. It involves two residual graph convolutional neural networks. GemSpot [17] is useful for the robust identification of ligand poses and drug discovery efforts through Cryo-EM. This tool aids in the correct interpretation and modeling of ligand densities and will greatly help drug discovery efforts based on Cryo-EM.

### 1.3 Introduction to Our Pipeline

To support our research, we introduce the EMD-Ligand Dataset, the first proteinligand binding database created from EMDataResource and Protein Data Bank (PDB). We evaluate current PLBP methods/models (with modifications so that they can apply to large protein complexes) on this new database, as well as testing these methods against the conventionally used, publicly-accessible protein-ligand binding databases which will be introduced in Section 2.1.1. Furthermore, we demonstrate how integrating PLBP methods with DeepTracer [18], a fully-automated deep learning-based method for de novo multichain protein complex structure modeling based on Cryo-EM maps, can create a fast and automated pipeline to directly predict ligand binding sites from Cryo-EM maps. Additionally, we show how extending this pipeline with Pocket2Mol [19], a deep generative model for efficient molecular sampling based on 3D protein pockets, can generate high-quality drug-like molecules without any prior design input.

In Section 2, we begin with a comprehensive comparative study of three popular PLBP methods (PUResNet, Kalasanty, DeepSurf) across multiple public datasets. Through our independent study, we aim to fairly assess these methods and use the best methods in our PLBP study on Cryo-EM maps. We found that there is no single method that consistently outperforms the others, so we evaluated all three methods on the EMD-Ligand Dataset. Next, we present the DeepTracer Protein Ligand Binding Prediction pipeline (DeepTracer-PLBP) and demonstrate its effectiveness. We also show the extended DeepTracer Drug-Like Molecules Generator pipeline (DeepTracer-DLMGen). In Section 3, we present the detailed evaluation results of these PLBP methods, as well as the evaluation of DeepTracer-DLMGen generated drug-like molecules based on the metrics that are widely used to evaluate the quality of generated drug candidates. Finally, in Section 4, we conclude with a discussion and our conclusions covering the case-proven successes and failures of utilizing DeepTracer in a pipeline to predict ligand binding sites and generate de novo drug-like ligands from Cryo-EM maps. The appendix contains the sample list of EMD maps whose ligand binding sites are correctly predicted by our DeepTracer-PLBP pipeline.

We showcase the designed high-level workflow of our method in Figure 1, which integrates the DeepTracer pipeline to predict ligand binding sites and generate druglike ligands directly from Cryo-EM maps. Our multi-step approach starts with an input Cryo-EM map and utilizes the DeepTracer pipeline to generate the protein’s backbone structure. This is then followed by the prediction of ligand binding sites using one of the PLBP methods. Finally, both the predicted protein structure and the predicted binding site are fed to Pocket2Mol to generate drug-like ligands.

**Fig. 1.**
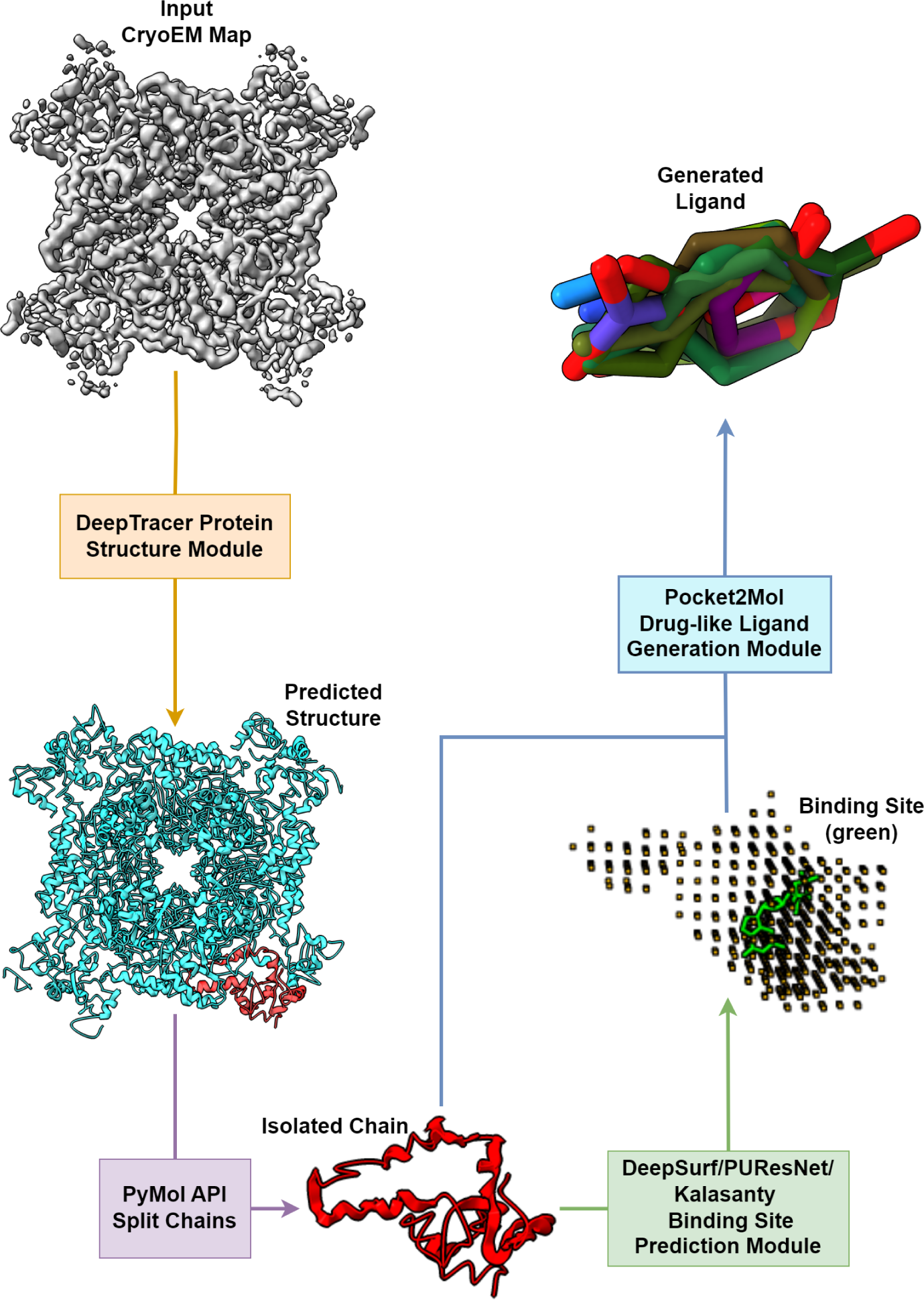
DeepTracer Ligand Binding Site Prediction and the Drug-like Molecules Generator pipeline. (created in ChimeraX).

## 2 Our Pipeline

### 2.1 Comparative Studies of State-Of-The-Art PLBP Methods

To begin, we introduce the popular datasets used for protein-ligand binding studies, followed by a discussion of our evaluation metrics. We introduce the three state-of-the-art methods that we are evaluating, namely Kalasanty, PUResNet, and DeepSurf. After that, we introduce our newly created EMD-Ligand Dataset which provides the feasibility to do the benchmark experiment. Further, integrating with DeepTracer will be introduced. Last, it is the case proven introduction of the integration with Pocket2Mol drug-like ligand generator.

#### 2.1.1 Existing Datasets for Protein-Ligand Binding Study

Examining the datasets used in training and evaluating PLBP algorithms is very helpful for this study. The PDB provides the 3D structures of many proteins and their ligands, and it is a widely used resource in structural biology. The BioLip [20], ScPDB [4], and BindingMOAD [21] databases provide additional curated data for studying ligand-protein interactions. The coach420 [22] and BU48 [23] datasets are independent test sets used to evaluate the performance of machine learning models in predicting protein-ligand binding sites. The core set within PDBBind [5] contains a diverse set of high-quality protein-ligand complexes, which serve as an essential benchmark dataset for different scoring functions.

The PDB database stores the primary information of protein structures, including coordinate files for all deposited biological molecules. These files list the atoms in each protein and their 3D location space. The basic syntax and format comprise category and attribute names. These entities describe the chemistry and identity of the molecules of a chemically distinct part of a structure.

PDBBind, the latest version updated in 2020 and a new 2021 version on the way, contains information for 23,496 protein-ligand binding interactions. A series of computer programs were implemented to screen the entire PDB database to identify four major types of molecular interactions: protein-small ligand, nucleic acid-small ligand, protein-nucleic acid, and protein-protein interactions [5]. It contains a special subset [24], called the “core set”, with high-quality protein-ligand complexes with remarkable diversity in structure as well as binding data. This subset is used in the popular Comparative Assessment of Scoring Function (CASF) evaluation. It is composed of 285 protein-ligand complexes in 57 clusters [24].

BioLip [20], updated weekly, is a curated database of ligand-protein binding interactions that have biological relevance. After the automated process for determining a ligand’s biological relevance is completed, a thorough manual review is performed to correct any errors. The structural data is primarily obtained from the PDB while biological insights are obtained from literature and other databases. Eventually, the manual check is performed and possible false-positive entries are verified by reading the original literature and consulting other databases that ensure the completeness and high quality of BioLip. The latest version (2023) has a total of 833,327 entries.

The ScPDB [4] database is a comprehensive and up-to-date selection of druggable binding sites of the Protein Data Bank. Druggable sites are defined as complexes between a protein and a pharmacological ligand. The database provides the all-atom description of the protein, the associated ligand, the binding site, and the binding mode of the protein-ligand interaction. It is continuously updated. The ScPDB release 2017 version is the most current release and it archives 16,034 entries, 4,782 proteins and 6,326 ligands [4]. Its features include identifying binding sites, correcting structures, annotating protein function, noting ligand properties, and characterizing their binding mode. Each entry also contains a transformed 3D MOL2 file of the structure of each protein-ligand complex [4].

BindingMOAD [21] contains very high-quality ligand-protein binding, a collection of high-quality holo-crystal structures resolved higher than 2.5Å or better. Currently MOAD release 2020 has 41,409 protein-ligand complexes and 15,223 entries of binding data. Coach420 [25] is an independent test dataset which consists of 420 protein structure with known ligands. We will use the dataset provided by PUResNet which utlizes the Coach420 dataset but removed 122 protein structures since they were present in PUResNet’s training dataset. Therefore, 298 protein structures with ligands were selected. BU48 [25] is another test set consisting of 48 proteins with known ligands.

#### 2.1.2 Evaluation Metrics

There are two standard metrics utilized for the evaluation: D_CC_ [26] and DVO [1],[26]. D_CC_ is the measured distance between the predicted and actual center of the binding pocket. It is typically used to describe a success rate for a method, i.e., the fraction of sites below the given D_CC_ threshold. DVO (discretized volume overlap) is a metric that compares the shapes of the predicted and actual pockets. It is the volume of the intersection of the predicted and the actual segmentations, divided by volume of their union. These two metrics complement one another, highlighting different aspects of prediction quality – correct location (D_CC_) and shape (DVO). An O_PL_ matrix [26] indicates the intersection, at the atomic level, between the real and predicted binding sites divided by their union. It differs with the above distance-based metrics by also considering the shape of the binding sites, since it expresses a normalized spatial overlap between the predicted and the actual location of the binding pocket. F_1_-score is a standard metric for detection tasks. It combines precision (positive predictive value) and recall (also called sensitivity) using the harmonic mean. Success rate (SR) is defined as the total number of successful predictions for all proteins divided by the corresponding total number of existing sites.

#### 2.1.3 PLBP Methods for Evaluation

All three methods use ScPDB for model training. K-fold cross-validation is used to tune hyperparameters and the final model was trained using the full dataset for best performance [27]. Kalasanty presents a deep learning-based approach for finding binding pockets, which was inspired by semantic image segmentation instead of classification. It used a 3D U-Net model structure [28]. The input to the model is the 3D coordinates of a protein, which is internally represented as a grid. Output is the corresponding ligandability scores of each vortex of the grid.

PUResNet improves Kalasanty by using a novel training data cleaning process based on structural similarity and a new model structure [2]. The data cleaning process groups protein structures according to the UniProtID and the Tanimoto index calculated for the high similarity of the protein structure in the cluster, followed by manual inspection using PyMol. Finally, 5,020 protein structures, out of 16,034, were selected from ScPDB dataset for training. Its model structure is derived from the concept of U-Net and ResNet by combining the convolutional segmentation of biomedical images with a skip connection between the blocks.

DeepSurf is a novel computational method for the prediction of potential binding sites [3]. It adopts ResNet architecture as its network backbone, which effectively combines the learning capabilities of advanced CNN architecture with a surface-based representation of the protein structure [29]. Two types of model architecture were investigated. One is the standard 3D-ResNet, and the other is a novel bottleneck 3D-LDS-ResNet, which shows better accuracy with a much smaller model size. We evaluated three methods, namely PUResNet, Kalasanty, and DeepSurf, on the Coach420 and BU48 sets as well as the PDBbind Core-Set (CASF-2016). The evaluation results are shown in Section 3.

### 2.2 Creation of Our EMD-Ligand Dataset

Cryo-EM technology has gained significant popularity since its introduction and an additional explosion in popularity following the advent of the “Resolution Revolution” that began in 2014, leading to an increase in experimentally-determined proteins in PDB banks using Cryo-EM. It produces 3D electron density maps of macromolecules and can explain the shape and interactions of protein complexes which forms the basis for modeling various pathologies, as well as assisting therapeutic drug design. Therefore, studying protein-ligand interactions based on Cryo-EM maps has become crucial. While ScPDB is the most popular database for protein ligand binding site prediction research, it mostly contains proteins with a few thousand atoms or fewer and higher resolution structures determined by x-ray crystallography (*<*2.5Å) and NMR. In contrast, protein structures found within the Cryo-EM database are typically large complexes with many chains, contain significantly more atoms, on the range of tens of thousands of atoms, and lower resolution (*>*3.0Å). Thus, available protein-ligand binding site prediction methods have not been utilized to test or report results on proteins with Cryo-EM maps.

To identify protein structures with both Cryo-EM maps and ligand-binding information, we used a database cross-query. Firstly, we queried the comprehensive data source for Cryo-EM maps in EMDataResource and found 24,505 Cryo-EM maps corresponding to 13,944 PDB coordinate entries in the PDB database. We downloaded the full PDB ID list (13,857) with detailed information, including 3D structures for proteins and ligands. We searched each of these 13,857 PDB IDs of the Cryo-EM maps dataset in all publicly available protein-ligand binding datasets and found no matches in ScPDB, no matches in MOAD dataset, 19 matches in PDBbind, and 2,878 matches in the BioLip dataset.

Since many Cryo-EM proteins are large protein complexes with numerous chains, the BioLip dataset was used to provide information on each protein and its respective ligand(s) (which were split according to chain). Chains with found ligands were saved as individual pdb files, along with their corresponding ligands. In total, the 2,878 matched proteins corresponded to 41,542 protein chain files and 46,538 corresponding ligands. To focus on our study’s objective, we removed all RNA, DNA, peptide, and ion (Zn, Mg, etc.) ligands, leaving 24,978 remaining ligands, corresponding to 1,621 proteins and 10,463 protein chains. Table 1 gives a breakdown of these statistics ranked by Cryo-EM map resolution, with 66% of Cryo-EM maps having resolutions lower than 3.0Å. This dataset forms the basis for our subsequent studies.

**Table 1.** Cryo-EM maps groups by resolution.

| Resolution(Å) | Number of Proteins | Number of Protein Chains | Number of Ligands |
| --- | --- | --- | --- |
| All | 1621 | 10463 | 24978 |
| 0.5–3.0 | 400 | 2689 | 8318 |
| 3.0–3.5 | 562 | 3529 | 8496 |
| 3.5–4.0 | 358 | 2807 | 5793 |
| 4.0–4.5 | 149 | 613 | 1207 |

### 2.3 Benchmark on EMD-Ligand Dataset Using the Ground-Truth Protein Structure

DeepSurf is a surface-based method that can directly predict binding sites on relatively large protein complexes with multiple chains but it has a hard-coded limit on the size of the proteins in terms of the number of surface points. On the other hand, Kalasanty and PUResNet assume that the protein structure is within a 70Å x 70Å x 70Å box, with all residues outside the box being ignored. These constraints make these methods either fail or perform very poorly on larger proteins with many chains, so we split the proteins according to their respective chains and apply the PLBP method to each individual chain. We evaluated Kalasanty, PUResNet, and Deep-Surf methods on our new EMD-Ligand Dataset. Unlike ScPDB and other datasets, which have only one ligand per protein, our dataset has multiple ligands per protein chain, with an average of nearly 2.5 ligands per protein chain. This means that the ligand-generation methods would generate multiple hypotheses as well, so we needed to modify our scoring algorithm to account for that.

We use the ground-truth protein chains (3D structures experimentally-determined from Cryo-EM maps) as input for all methods to predict ligand binding sites. Their evaluation results are given in Section 3. As an illustration, Figure 2A shows the structure of EMD-5127 chain G (corresponding PDB ID: 3iyd) with a reference binding ligand (CMP) shows the DeepSurf-predicted ligand binding site (the pink dotted line on the right) with respect to potential interacting protein residues. Figure 2B shows the DeepTracer-predicted protein structure EMD-21391 chain M (corresponding PDB ID: 6vv5) with the reference ligand PAM showing the PUResNet predicted ligand binding site. As we can see, the binding site surrounds the ligand, and the center of the predicted pocket is close (*<*4.0Å) to the center of the ligand.

**Fig. 2.**
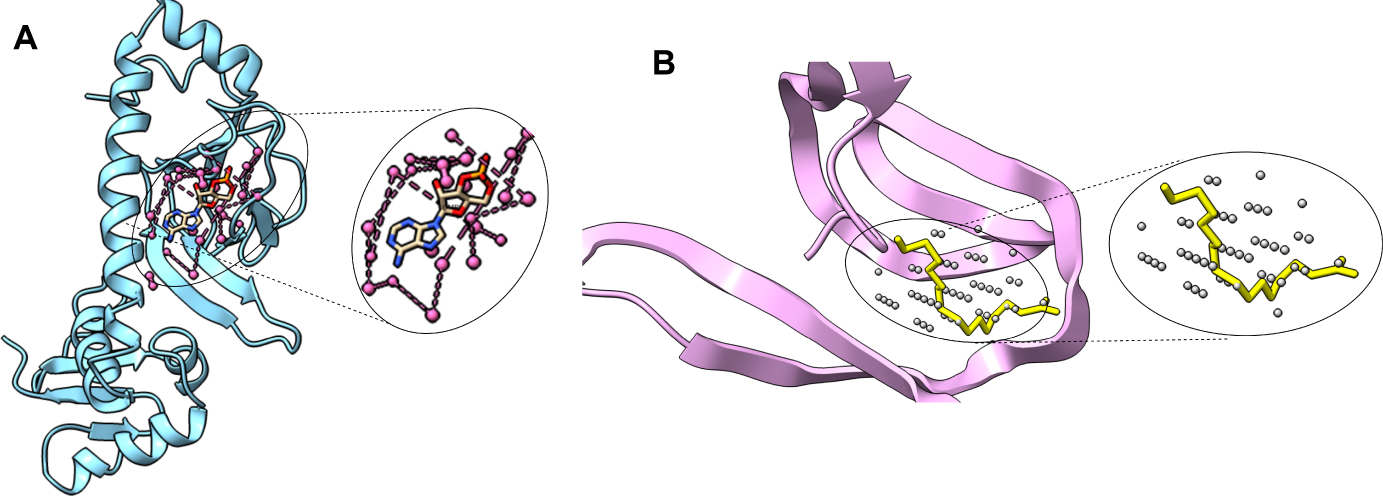
A) The protein-ligand binding site prediction using ground truth protein structure. (PDB ID: 3iyd), chain G (EMD-5127) with its ligand CMP. The purple dotted line represents the predicted binding site by DeepSurf; CMP is depicted inside in the ground truth state. This is a correct prediction as the D_CC_ is less than 4.0Å and shows that the predicted site overlaps with the ligand (created in ChimeraX). B) The protein-ligand binding site prediction using DeepTracer-predicted protein structure. EMD-21391(PDB ID: 6vv5), chain-M with the PUResNet predicted pocket and the ligand PAM (yellow). Grey dots represent the PUResNet-predicted binding pocket (created in ChimeraX).

### 2.4 Integration with DeepTracer Protein Structure Prediction Pipeline

DeepTracer is a powerful deep learning-based method and service for protein structure modeling using Cryo-EM maps [18]. It uses a 3D convolutional neural network that allows for fast and accurate structure predictions. Compared to other state-of-the-art methods such as Phenix [30], Rosetta [31] [32] and MAINMAST [32], applied on a set of coronavirus-related Cryo-EM maps, there is on average 30% more residue coverage when compared to Phenix with an average RMSD improvement of 0.11Å; applied on a set of nine Cryo-EM maps, RMSD improvements range from 1.37Å to 0.85Å; yielding 57% more coverage than MAINMAST. To extend the functionality of this method, we integrate DeepTracer with two other PLBP methods, PUResNet and DeepSurf, which have demonstrated promising results on the EMD-Ligand Dataset. In doing so, we have built an end-to-end pipeline (DeepTracer-PLBP) that can directly predict protein-ligand binding sites from Cryo-EM maps without requiring experimentally-determined protein structures.

As shown in Figure 3, we present another example of this pipeline using the protein Holo-bacterioferritin-form-I (EMD-9913) in 3A the ground-truth state as well as 3B the DeepTracer-predicted structure of chain V. In 3C, we show an overlay of the ground-truth state and the DeepTracer-predicted structure with its corresponding ligands, Fe^2+^ (red) and HEM (cyan). Using PUResNet, the predicted ligand binding pocket (as shown with grey dots), which overlaps with the true ligand, is validated based on the distance between the center of the binding pocket and the center of the ligand itself with a distance *<*4.0Å.

**Fig. 3.**
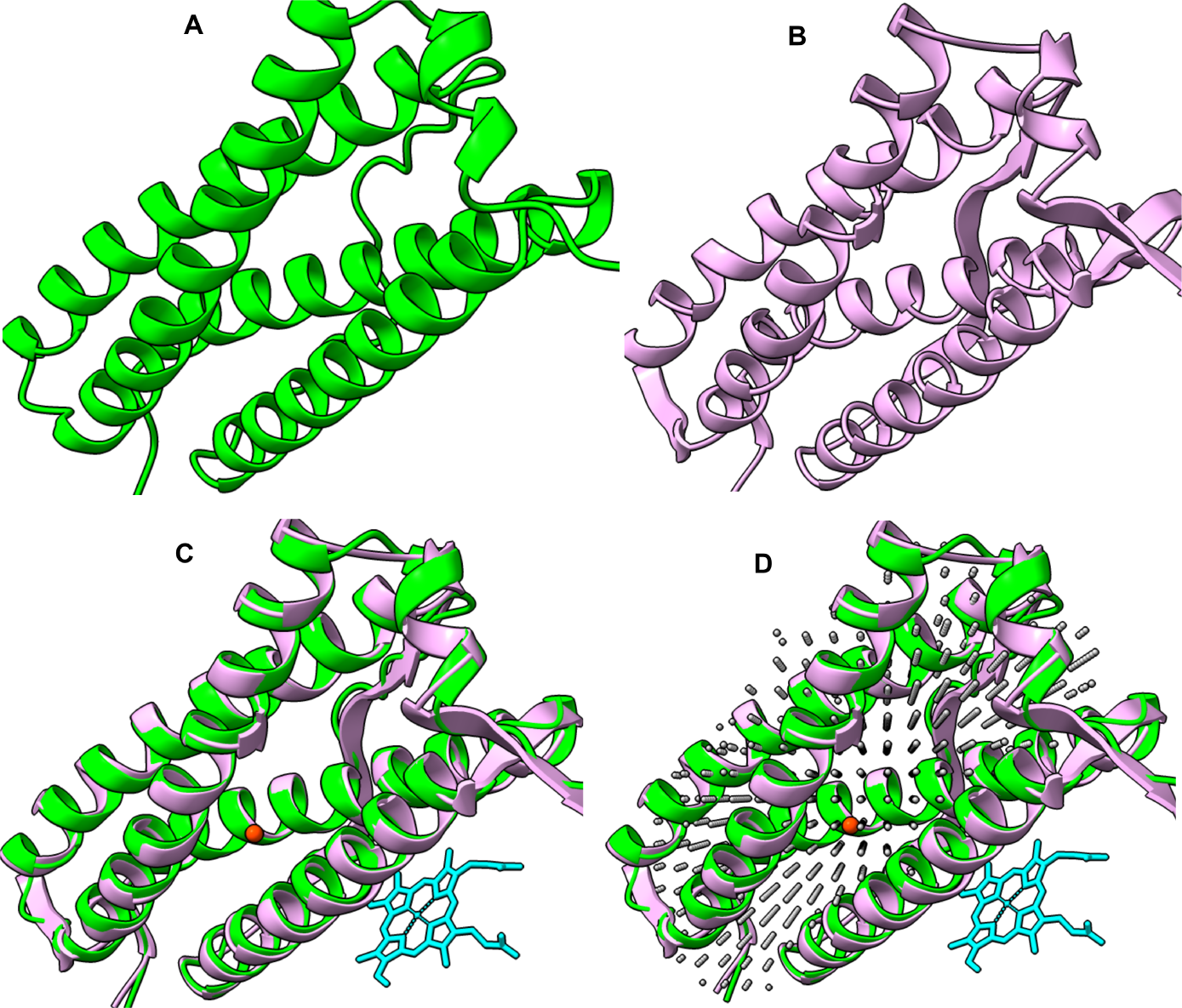
The integration of ligand binding site prediction with DeepTracer. A) The ground truth state of the protein structure (PDB ID: 6K43, chain N). B) The DeepTracer-predicted protein structure of EMD-9913, chain V. C) The protein structure with two ligands, orange dot (Fe^2+^), light blue (HEM). D: the gray dots represent the ligand binding site predicted by PUResNet, validated based on a D_CC_ value of less than 4.0Å to the ligands.

To demonstrate the effectiveness of DeepTracer-PLBP, we used the EMD-Ligand Dataset. However, we needed to make a small modification to our scoring algorithm since the predicted protein structures by DeepTracer may have a different number of chains compared to the ground-truth structures, as the chain IDs may not correspond to the same section of the protein of interest. Therefore, we compare each ligand prediction to all reference ligands of the full protein complex, not just one chain.

As depicted in Figure 4, the pipeline begins by taking a de novo multichain protein complex with a Cryo-EM map as input to DeepTracer, which predicts the backbone structure. Next PyMol is used to split all predicted protein chains into individual units. Lastly, each chain is fed into either PUResNet or DeepSurf to predict possible ligand-binding sites.

**Fig. 4.**
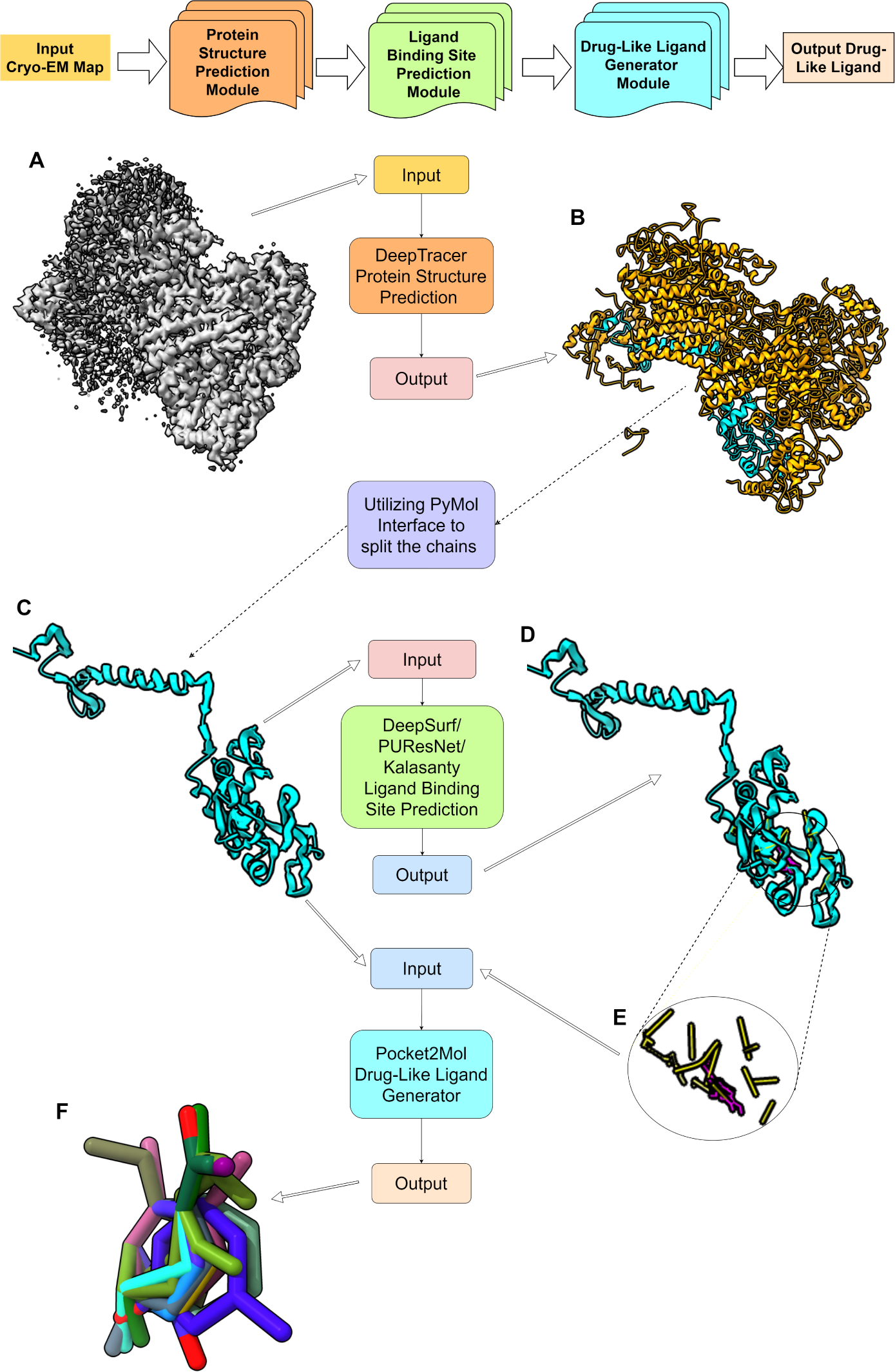
The high-level workflow pipeline of DeepTracer-PLBP and DeepTracer-DLMGen from an input Cryo-EM map. A: Input Cryo-EM map EMD4165. B: DeepTracer-predicted protein structure. C: Chain A of the DeepTracer-predicted protein structure. D and E: DeepSurf predicted pocket (yellow) with the ligand (purple) on chain A. F: seven generated drug-like ligands (overlapped in different colors).

### 2.5 DeepTracer Drug-like Ligand Generator

The ligand binding site prediction capability of DeepTracer can be utilized for in silico drug design through integration with the Pocket2Mol pipeline [19]. Pocket2Mol is an E(3)-equivariant generative network composed of a new GNN that captures both spatial and bonding relationships between atoms of the binding pocket and a new efficient algorithm that samples new drug candidates conditioned on the pocket representations from a tractable distribution without relying on conventional MCMC algorithms [33]. According to Peng et al., evaluation of ligand-generation performance of Pocket2Mol is completed through comparison to other existing, state-of-the-art methods such as a CVAE-base model [19] and another auto-regressive generative model [33]. The result shows that Pocket2Mol can sample drug candidates with more accurate binding properties including binding affinity, drug-likeness, and synthetic accessibility.

To demonstrate that DeepTracer-PLBP pipeline can be extended to generate drug-like molecules, we appended the Pocket2Mol pipeline to it. As shown in Figure 4 below, the DeepTracer-predicted protein structure is provided to PUResNet to predict possible ligand-binding sites. The coordinates of the center of the predicted binding site and the predicted protein structures are then provided to Pocket2Mol which uses the chemical and geometric attributes of the binding pockets to generate candidate ligands for the predicted binding site.

Figure 5 demonstrates the capability of the DeepTracer-DLMGen pipeline. Panel A depicts the predicted structure of Alternative complex III (PDB ID: 6F0K - EMD-4165) chain A. Panel B demonstrates several Pocket2Mol-generated drug-like molecules (white) located in the ligand binding pocket predicted by DeepSurf. This pipeline produces various DLM candidates, though for space considerations only a few are plotted in the figure. Panel C and D show the DeepTracer-predicted structure, PUResNet-predicted binding site, and the Pocket2Mol-generated DLMs of Human IMPDH2 (PDB ID: 6UAJ - EMD-20707) chain R.

**Fig. 5.**
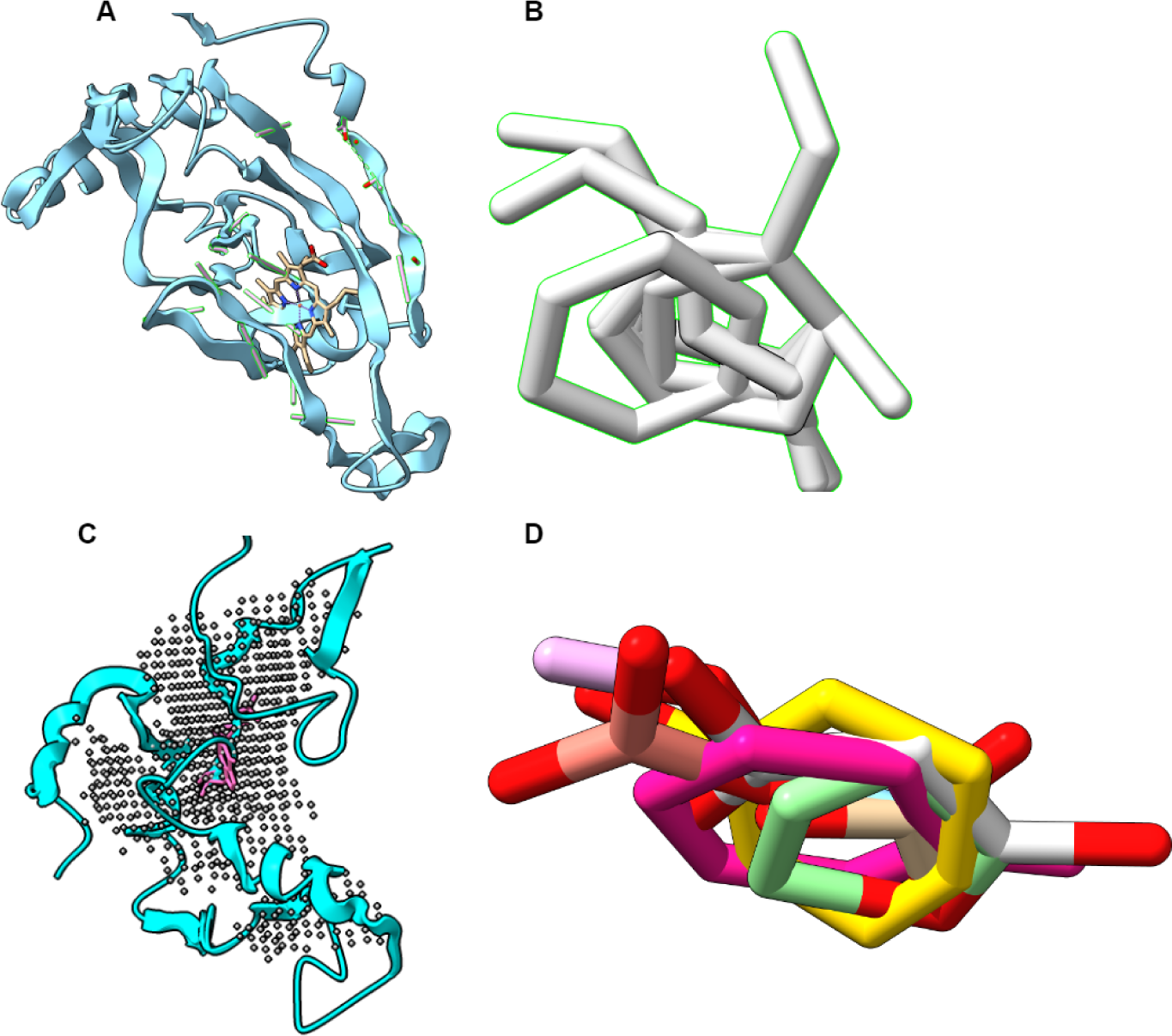
DeepTracer-BindPRED and DLMGen of Alternative complex III (PDB ID: 6F0K - EMD- 4165) chain A and Human IMPDH2 (PDB ID: 6UAJ – EMD-20707). A) The DeepTracer-predicted structure of Alternative complex III with the DeepSurf-predicted ligand binding site bound with HEC (represented by sticks). B) The DLMs generated by Pocket2Mol (white). C) DeepTracer-predicted structure of Human IMPDH2, chain R with the binding site predicted by PUResNet (shown in gray dots) and bound GTP (purple). D) The DLMs generated by Pocket2Mol.

## 3 Experiments and Results

### 3.1 Evaluation of PLBP Methods’ Performance on the Traditional Protein-Ligand Binding Databases

We evaluate the effectiveness of PUResNet, Kalasanty, and DeepSurf on the Coach420, BU48, and PDBbind Core-Set (CASF-2016) datasets to provide a comprehensive comparison of their binding-site prediction capabilities. It is important to note that PUResNet expresses its predicted binding site in terms of the grid in the pocket or cavity where the ligand is located while Kalasanty expresses its predicted binding sites in terms of the atoms and/or residues that are near the predicted site (using a distance cut-off threshold of 4.0Å). Therefore, the predicted sites of these two methods cannot be directly compared. To ensure a fair comparison, we modifed the Kalasanty algorithm to express its prediction in terms of the pocket or cavity as well. Figure 6 (left) demonstrates that the PUResNet prediction produces a slightly better D_CC_ curve on the Coach420 dataset. However, the DeepSurf method utilizes surface-point based prediction and thus can only express binding sites in terms of the atoms/residues on the protein surface. To compare DeepSurf with Kalasanty and PUResNet, we modifed the PUResNet method to output atoms and/or residues close to its predicted binding sites in the exact same way as Kalasanty does. Figure 6 (right) shows the D_CC_ curves of predicted binding sites (not pockets/cavities) of these three methods on the Coach420 dataset.

**Fig. 6.**
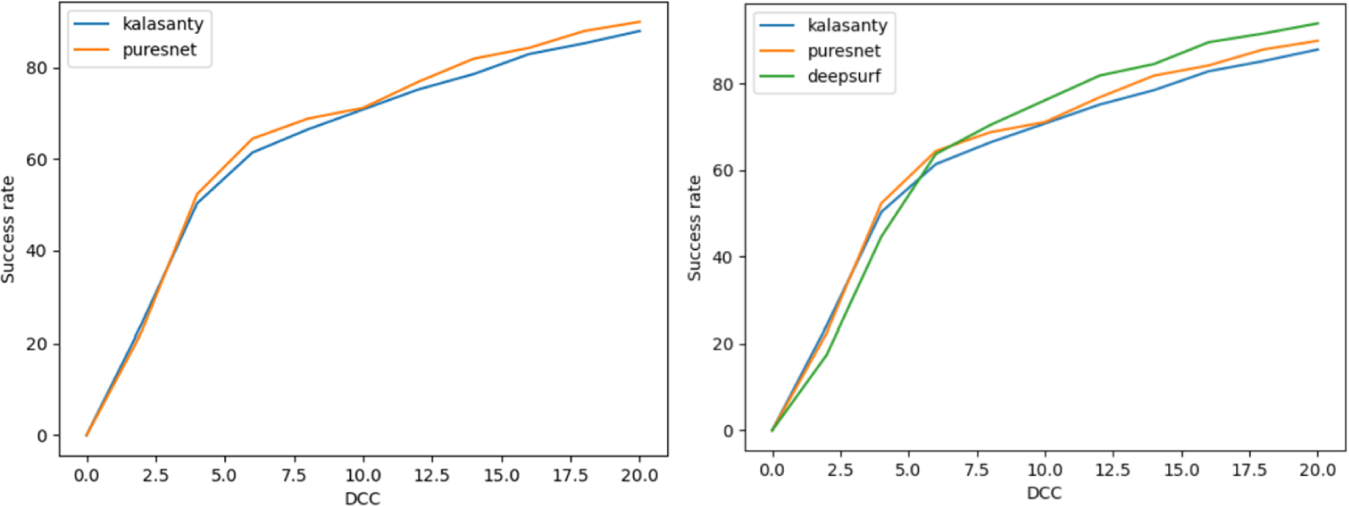
(left) D_CC_ performance of Kalasanty (cavity) and PUResNet on testing dataset Coach420. (right) D_CC_ performance of Kalasanty, PUResNet and DeepSurf on testing dataset Coach420, comparing predicted sites (residuals).

When D_CC_ falls below 4.0Å, it is considered a valid prediction. Using this criterion, we can calculate True Positive (TP), False Positive (FP), False Negative (FN), F_1_-score, and Success Rate (SR). Table 2 shows the results comparison of these three methods applied to three chosen test datasets: Coach420, BU48, and CASF-2016. For the Coach420 dataset, DeepSurf achieves the best F_1_-score and SR, while PUResNet is only slightly better than Kalasanty. For the BU48 dataset, DeepSurf and PUResNet both achieve a *∼*60% SR, and DeepSurf has the best F_1_-score due to significantly fewer FPs. However, Kalasanty performs the worst on this dataset, demonstrating a clear gap from PUResNet even though they perform similarly on the larger Coach420 dataset. On the PDBbind Core Set, Kalasanty performs significantly better than PUResNet. In summary, no one method clearly outperforms the other methods on all test sets.

**Table 2.** Comparison of Kalasanty and PUResNet on predicted binding sites on the Coach420 dataset. Performance comparison of predicted binding sites (represented by atoms/residues on the protein, not by pockets/cavities) on Coach420 set, containing 298 protein-ligand complexes. Evaluation results of each method on the BU48 dataset, containing 62 protein-ligand complexes. Evaluation results of each method on the PDBbind Core Set (CASF-2016), containing 285 protein-ligand complexes. (DeepSurf results are not included in last row due to unexpected results – see explanation above).

| Test Dataset | Model | TP | FP | FN | F<sub>1</sub>-score | SR | Declared SR<sup>1</sup> |
| --- | --- | --- | --- | --- | --- | --- | --- |
| Coach420 | Kalasanty | 119 | 174 | 25 | 0.54 | 40.0% | 51% |
|  | PURESNet | 122 | 175 | 19 | 0.56 | 40.9% | 53% |
|  | DeepSurf | 133 | 184 | 11 | 0.58 | 44.6% | N/A |
| BU48 | Kalasanty | 27 | 33 | 7 | 0.574 | 43.5% | 57% |
|  | PURESNet | 37 | 34 | 3 | 0.667 | 59.7% | 62% |
|  | DeepSurf | 36 | 27 | 2 | 0.713 | 58.1% | N/A |
| CASF-2016 | Kalasanty | 165 | 110 | 30 | 0.702 | 57.9% | N/A |
|  | PURESNet | 130 | 167 | 17 | 0.586 | 45.6% | N/A |

<sup>1</sup>from Peng et al, 2022

### 3.2 Evaluation of PLBP Methods’ Performance on the Novel EMD-Ligand Dataset

We utilize the ground-truth all-chain structures as input for all methods to predict ligand binding sites. The evaluation results are given in Table 3. The reference ligand information is given and the default D_CC_ threshold of 4.0Å is used. To understand the impact of Cryo-EM map resolution, we broke down the results into groups based on map resolution.

**Table 3.** Evaluation results of Kalasanty, DeepSurf, and PUResNet on the novel EMD-Ligand dataset. DeepSurf performance on resolution 4.0Å-4.5Å subset with different D_CC_ thresholds in scoring. Results of the original Kalasanty model vs. the adapted Kalasanty model (Trained on the 3.0Å-3.5Å subset and evaluated on the 3.5Å-4.0Å subset)

| Resolution (Å) | Model | $D_{CC}$ threshold | Pred-Ligand Number | FN | TP | FP | F<sub>1</sub>-score | SR (%) |
| --- | --- | --- | --- | --- | --- | --- | --- | --- |
| 0.5-3.0 | Kalasanty | 4.0Å | 1349 | 7939 | 378 | 971 | 0.078 | 4.60 |
|  | DeepSurf | 4.0Å | 3281 | 7798 | 517 | 2764 | 0.089 | 6.30 |
|  | PuresNet | 4.0Å | 1689 | 7718 | 597 | 1092 | 0.119 | 7.20 |
| 3.0-3.5 | Kalasanty | 4.0Å | 2729 | 7758 | 738 | 1991 | 0.132 | 8.70 |
|  | DeepSurf | 4.0Å | 3901 | 7663 | 830 | 3071 | 0.134 | 9.80 |
|  | PuresNet | 4.0Å | 3274 | 6944 | 1546 | 1728 | 0.263 | 18.30 |
| 3.5-4.0 | Original-Kalasanty | 4.0Å | 2620 | 5053 | 740 | 1880 | 0.176 | 12.80 |
|  | Adapted-Kalasanty | 4.0Å | 2160 | 4696 | 1097 | 1063 | 0.276 | 18.90 |
|  | DeepSurf | 4.0Å | 2588 | 5297 | 496 | 2092 | 0.118 | 8.60 |
|  | PuresNet | 4.0Å | 2798 | 4430 | 1362 | 1436 | 0.317 | 23.50 |
| 4.0-4.5 | Kalasanty | 4.0Å | 352 | 1117 | 90 | 262 | 0.116 | 7.50 |
|  | DeepSurf | 4.0Å | 1250 | 942 | 264 | 986 | 0.215 | 21.90 |
|  | DeepSurf | 5.0Å | – | 891 | 314 | 936 | 0.256 | 26.10 |
|  | DeepSurf | 6.0Å | – | 852 | 351 | 899 | 0.286 | 29.40 |
|  | PuresNet | 4.0Å | 485 | 1020 | 187 | 298 | 0.221 | 15.50 |

**Note for Table 3:** We did the above experiments for each resolution subset. The results prove promising for the continued research on our EMD-Ligand Dataset. There is evidence that the improving success rate from 12.8% to 18.9% following 3.0Å-3.5Å subset training and evaluation on the 3.5Å-4.0Å group.

As seen in Table 3, PUResNet performs the best overall (SR=16.1%) compared to DeepSurf (SR=15.6%) and Kalasanty (SR=6.4%). It is worth noting that all methods show large variations across different resolution groups, as expected. The highest resolution group (0.5Å-3.0Å) had the worst accuracy among all resolutions, which is surprising, especially considering that all these methods were trained using the ScPDB database in which all protein images are in that same resolution range. PUResNet had the best accuracy for all resolutions excluding the poor performance noted in the 4.0Å-4.5Å group in which DeepSurf performed better (SR=21.9%). PUResNet showed significantly better performance than the other two methods on the 3.5Å-4.0Å resolution group (SR=23.5%). Strikingly, the performances of all tested methods on this dataset, when compared to the non-Cryo-EM maps protein/ligand dataset reported in Section 3.1 (Table 2), are significantly worse. This clearly demonstrates the fragility of these PLBP methods across different datasets, though the reasons are unclear. To improve this methodology, more research is required.

The scoring algorithm used in our evaluation considers multiple hypotheses since the EMD-Ligand Dataset has multiple ligands per protein chain. However, all methods still produced many false negatives. We believe that this is the primary cause of their poor performance compared to the non-Cryo-EM maps protein/ligand dataset reported in Table 2. This may be due to the fact that all three examined methods were trained on the ScPDB database, which is very different from the EMD-Ligand Dataset in terms of map resolution and protein size. We hypothesize that adding more diverse training data into these methods will greatly increase their performance. To test this, we conducted a preliminary using Kalasanty. We aimed to refine or adapt the original Kalasanty model using protein-ligand data from the resolution 3.0Å-3.5Å subset and evaluate the adapted model on the 3.5Å-4.0Å subset. The results are given in Table 3. It is promising to observe a 6.1% increase in SR for the adapted Kalasanty model compared to the original model. This suggests that adding more diverse training data could improve the performance of these PLBP methods.

### 3.3 Evaluation of De Novo Drug-Like Molecule Generation

Several metrics are widely used to evaluate the quality of generated drug candidates [25]. The Vina score estimates the binding affinity between the generated molecules and binding site, with higher affinity being calculated as the percentage of sites in which the generated molecules have higher or equal affinity to those in the test dataset [34]. The Quantitative Estimation of Drug-likeness (QED) metric measures how likely a molecule is to be a potential drug candidate based on several criterion [17]. The Synthetic Accessibility Score (SAscore) represents the difficulty of drug synthesis [35]. The LogP value represents the octanol-water partition coefficient and indicates ideal drug candidates based on values ranging from −0.4 to 5.6 [36]. Finally, Lipinski’s rule of five measures how many rules a ligand follows to be considered drug-like [37].

According to Roy et al., 2012, Pocket2Mol has shown better scores in all these metrics when tested on the CrossDocked dataset compared to other methods [25]. The generated molecules not only demonstrate higher binding affinities and more stable chemical properties, but also produce more realistic and accurate structures. In our pipeline, we use the same metrics (excluding Vina Score and High affinity, which are either unavailable or inapplicable) to evaluate the quality of generated molecules. Results for four binding sites are provided in Table 4, where higher scores indicate better results. The scores are averaged over all generated molecules for each site. Compared to Roy et al., 2012, our results are very competitive, though analysis was run on a different dataset and average over many proteins structures [25].

**Table 4.**
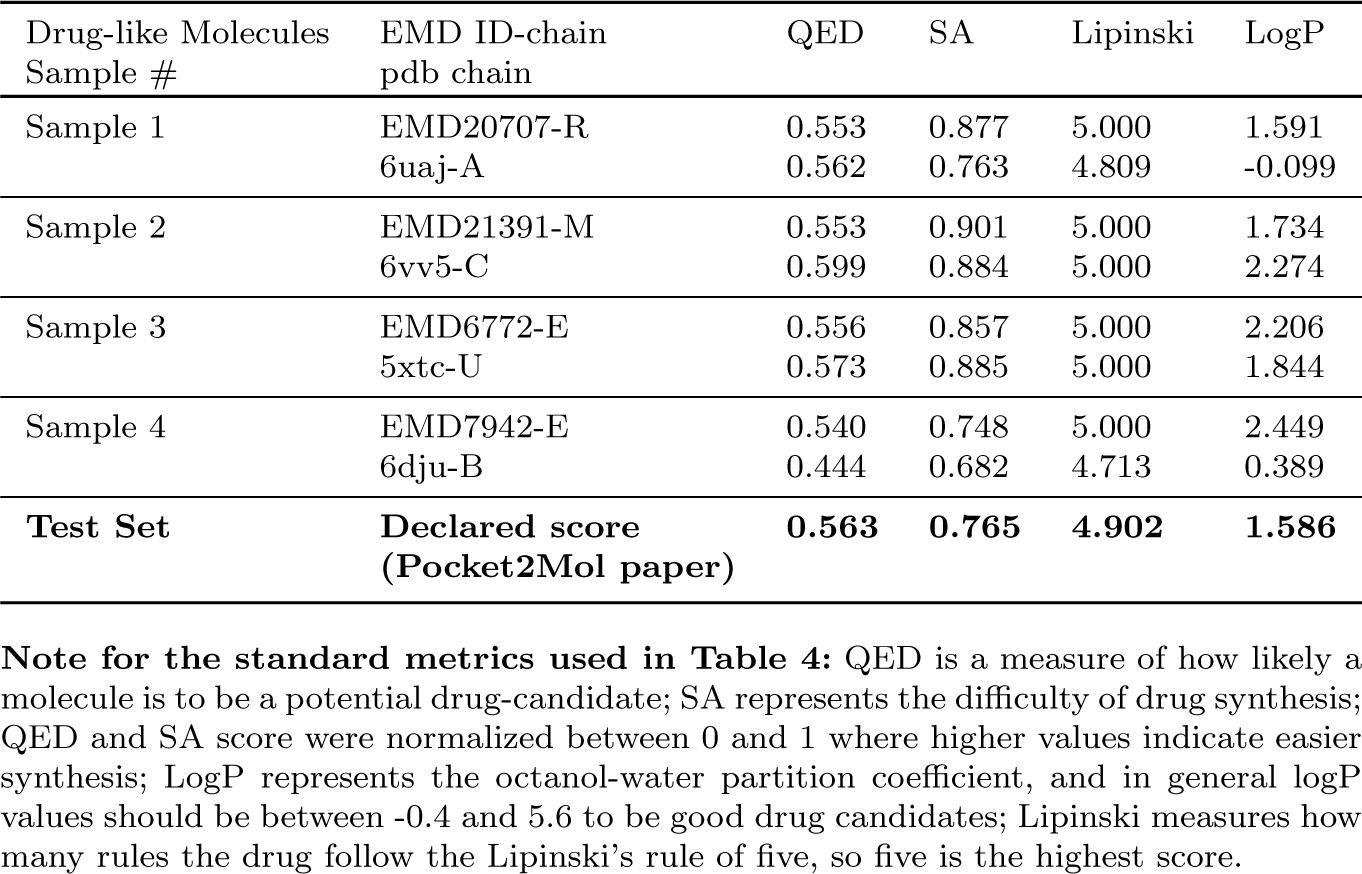
Evaluation of Pocket2Mol-generated molecules using DeepTracer-predicted protein structures and PUResNet/DeepSurf-predicted ligand-binding sites. A comparison of generated molecules to ground-truth protein structures.

To further compare these results, we also used the ground-truth protein structures and ligand-binding information as inputs for Pocket2Mol-based DLMGen. The results of this examination can be viewed in Table 4 below. For each sample, row 1 represents the DeepTracer-generated structure and the predicted ligand-binding site while the second row represents the corresponding ground-truth structure and ground-truth ligand-binding site. Using DeepTracer-predicted structures and ligand binding sites was as effective as using ground-truth structures, as shown by comparable validation criterion values. This demonstrates that DeepTracer-DLMGen is a valuable tool for drug design when applied to Cryo-EM maps.

## 4 Discussion and Conclusions

We conducted a comprehensive comparative study of three advanced deep learning-based methods for predicting protein-ligand binding sites (Kalasanty, PUResNet, and DeepSurf) on publicly available databases (Coach420, BU48 and CASF-2016). We then initiated the development of ligand-binding site prediction for proteins using Cryo-EM maps by introducing our DeepTracer-PLBP pipeline and created a novel EMD-Ligand Dataset specifically for this purpose. To our knowledge, this is the first database of its kind for predicting ligand binding in protein complexes with Cryo-EM maps. We evaluated the performance of these methods on this database and found that their accuracy is very limited compared to single-chain proteins with high-resolution structures determined by X-ray crystallography and NMR. This finding calls for more research efforts from the community to address this issue. Our preliminary experiments demonstrated that the addition of more training data, particularly proteins with Cryo-EM maps, can significantly improve their performance, even though there is still a large difference between them with their performance on the conventional datasets.

We also applied these methods to structures predicted by DeepTracer using Cryo-EM maps and successfully demonstrated that the integration of these methods with DeepTracer could create an efficient pipeline to predict ligand binding sites directly from Cryo-EM maps, eliminating the need for experimental methods and processes to determine protein structures, which can be costly and time-consuming.

We further extended the DeepTracer pipeline with Pocket2Mol to generate de novo drug-like molecules (DeepTracer-DLMGen) automatically and efficiently from Cryo-EM maps, which could be valuable for drug discovery and design. Our future work involves gathering more relevant datasets, particularly those with Cryo-EM maps, to train better models for this task. We plan to improve the existing deep learning models by introducing better model structures, such as attention mechanisms, better features, and model ensembles. Additionally, we believe that training models with DeepTracer-predicted protein structures, rather than only experimentally-determined structures, will significantly improve their performance. The success of the DeepTracer-PLBP could improve the accuracy and ease of drug discovery and generation and make the entire pipeline more impactful overall.

## Acknowledgments

We express our gratitude to the authors of DeepSurf and PUResNet for their generous help in answering our questions throughout this work. We would also like to extend our appreciation to the members of the DAIS team, including Jason Chen, Haowen Guan, and Natha Chiu, for their invaluable support. Additionally, we are grateful to Dr. Wooyoung Kim and Dr. Minglei Zhao for their insightful discussions and advice.

## Additional Information

### Code availability

The DeepTracer-PLBP pipeline will be available to use in the near future at https://deeptracer.uw.edu/

### How to cite this article

Lu, C. et al. Protein-Ligand Binding Site Prediction and de Novo Ligand Generation from Cryo-EM Maps (2023).

## Appendix

**The scoring algorithm’s pseudo-code:**

1. For each predicted pocket
2. Find the closest reference ligand (among all reference ligands for this protein chain)
3. If the distance D_CC_ *<*= 4.0Å, TP += 1
4. Else FP += 1
5. For each reference ligand
6. Find the closest pocket (among all predicted pockets for this protein chain)
7. If the distance D_CC_ *>*4.0Å, FN += 1
8. F_1_-score = 2 * TP / (2 * TP + FP + FN)
9. Success Rate = TP / Total ligands.

The following tables are sample lists of EMD maps (with chain) whose ligand binding sites are correctly predicted by our DeepTracer Ligand-Binding Site Prediction Pipeline (shown in Figure 6). Notation is as follows: “EMD-20707-R” means chain R of EMD map 20707. Reference “6uaj GTP A 1” is the ground truth ligand, 6uaj is the PDB id for EMD-20707. GTP is the ligand name, A is the chain id, “1” is the ligand index (6uaj chain A could have more than 1 ligand).

**Table 1.**
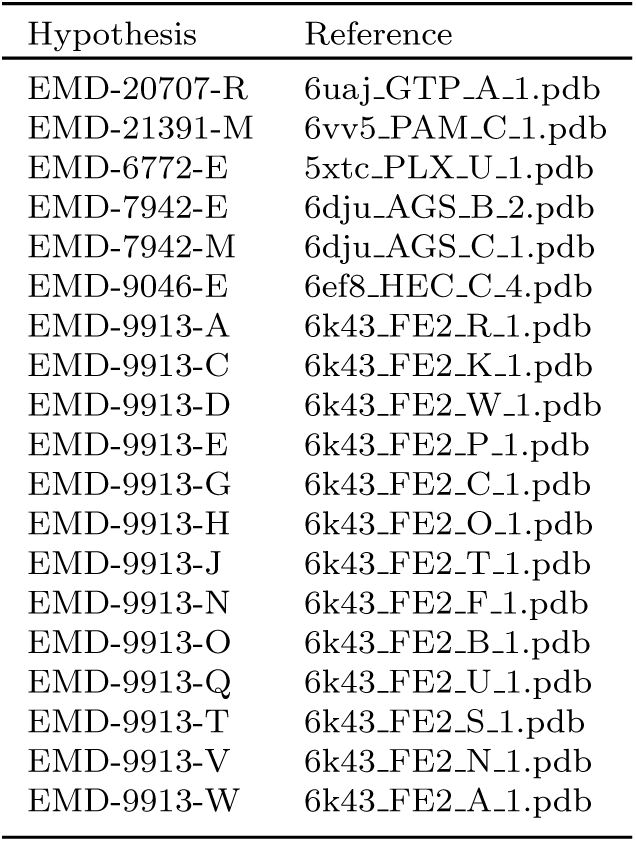
For resolution group 3.5Å–4.0Å, the correct prediction by PUResNet.

**Table 2.**
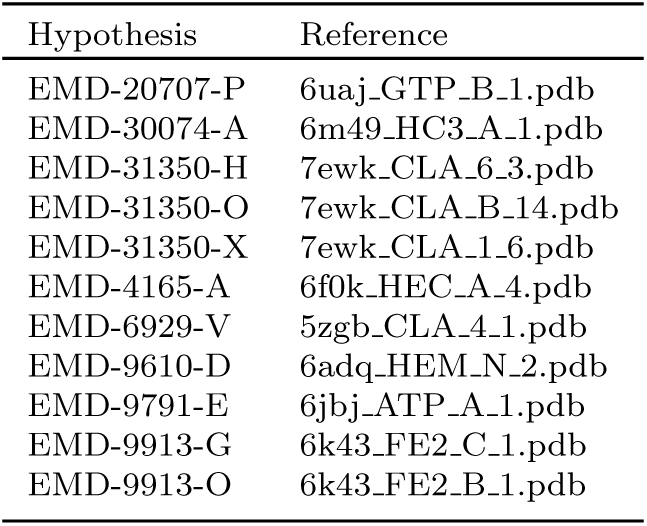
For resolution group 3.5Å–4.0Å, the correct predictions by DeepSurf.

**Table 3.**
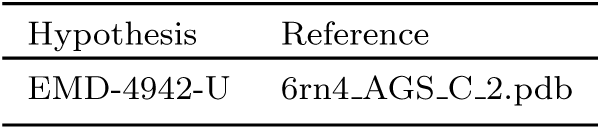
For resolution group 4.0Å–4.5Å, the correct predictions by PUResNet.

**Table 4.**
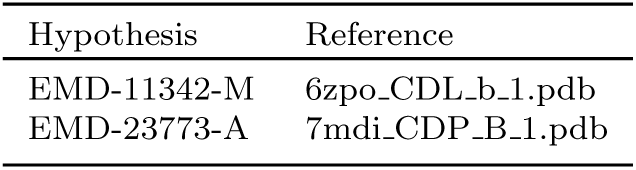
For resolution group 4.0Å–4.5Å, the correct predictions by DeepSurf.

**Table 5.**
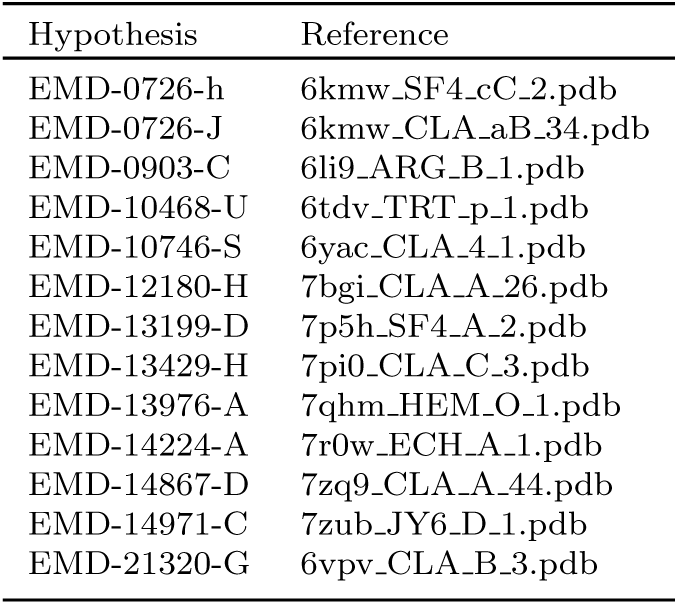
For resolution group 4.0Å–4.5Å, the correct predictions by PUResNet.

**Table 6.**
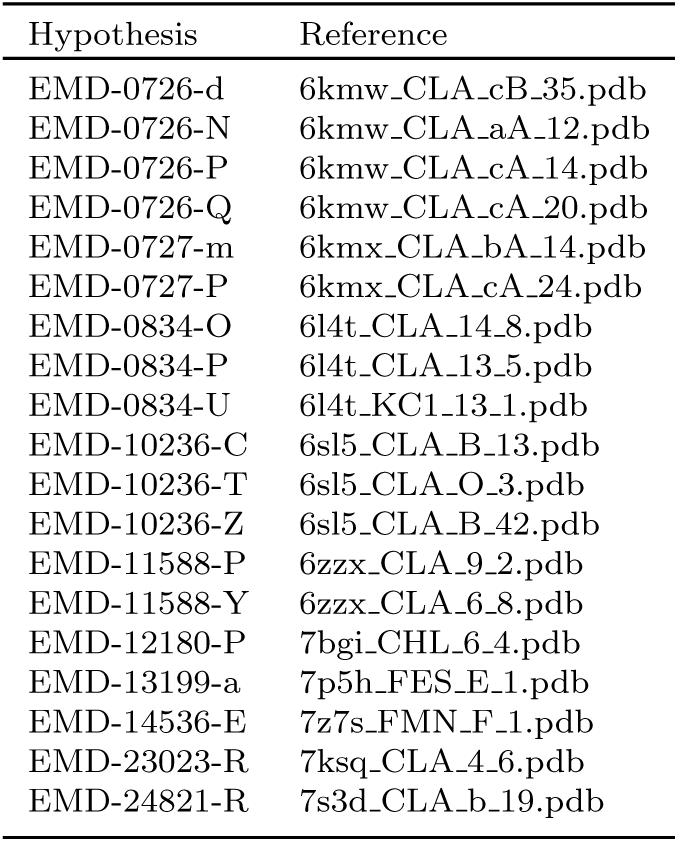
For resolution group 0.5Å–3.0Å, the correct prediction by DeepSurf.

**Table 7.**
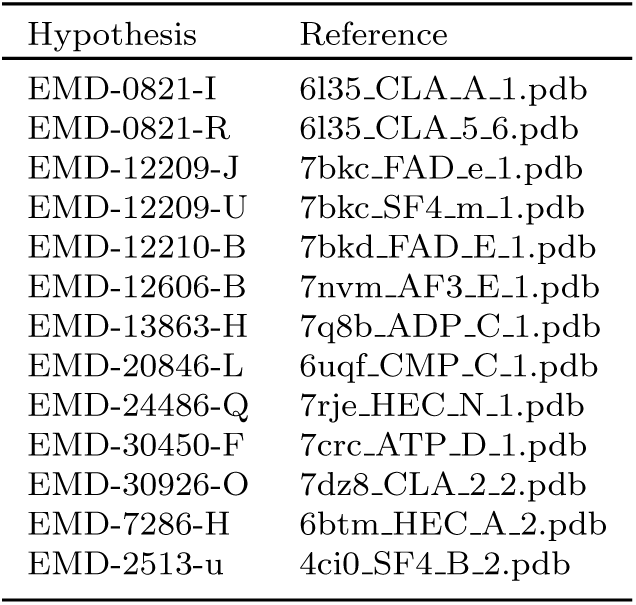
For resolution group 3.0Å – 3.5Å, the correct prediction by PUResNet.

**Table 8.**
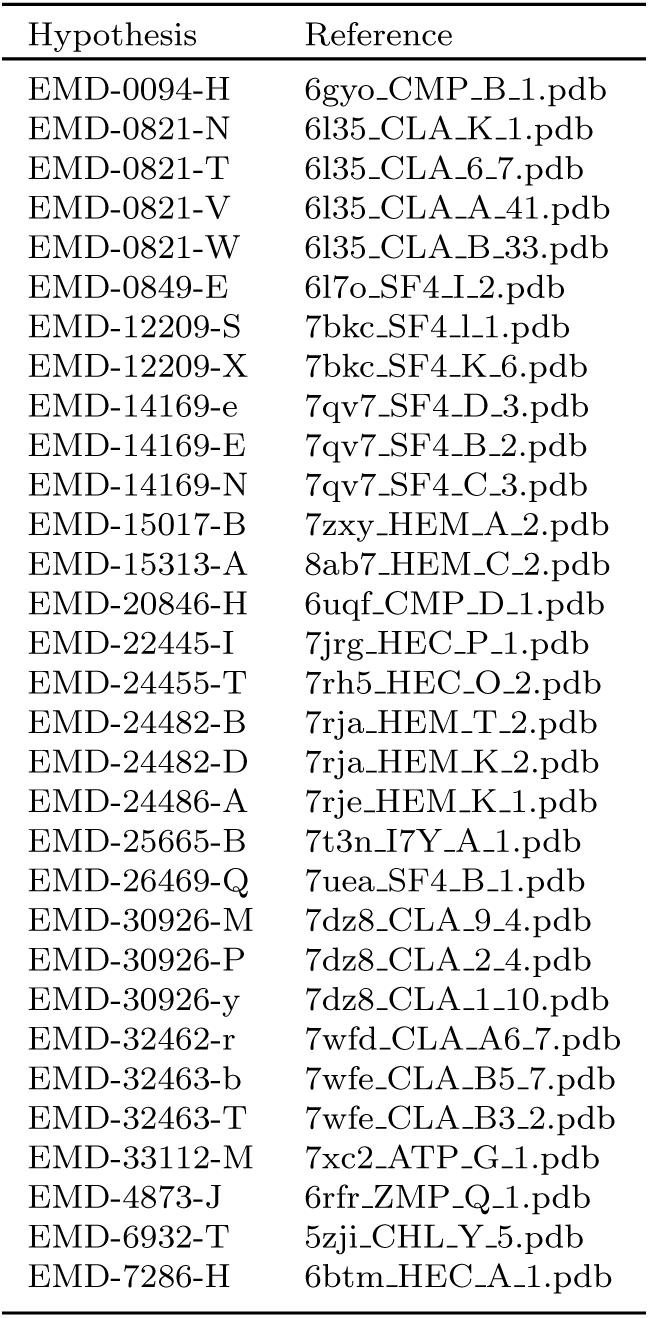
For resolution group 3.0Å – 3.5Å, the correct prediction by DeepSurf.

